# *Tahoe-100M*: A Giga-Scale Single-Cell Perturbation Atlas for Context-Dependent Gene Function and Cellular Modeling

**DOI:** 10.1101/2025.02.20.639398

**Authors:** Jesse Zhang, Airol A Ubas, Richard de Borja, Valentine Svensson, Nicole Thomas, Neha Thakar, Ian Lai, Aidan Winters, Umair Khan, Matthew G. Jones, John D. Thompson, Vuong Tran, Joseph Pangallo, Efthymia Papalexi, Ajay Sapre, Hoai Nguyen, Oliver Sanderson, Maria Nigos, Olivia Kaplan, Sarah Schroeder, Bryan Hariadi, Simone Marrujo, Crina Curca Alec Salvino, Guillermo Gallareta Olivares, Ryan Koehler, Gary Geiss, Alexander Rosenberg, Charles Roco, Daniele Merico, Nima Alidoust, Hani Goodarzi, Johnny Yu

**Author notes:** These authors contributed equally (Jesse Zhang, Airol A Ubas, Richard de Borja, Valentine Svensson, Nicole Thomas). These authors contributed equally (Neha Thakar, Ian Lai, Aidan Winters, Umair Khan, Matthew G. Jones). These authors contributed equally (John D. Thompson, Vuong Tran, Joseph Pangallo, Efthymia Papalexi, Ajay Sapre, Hoai Nguyen, Oliver Sanderson, Maria Nigos, Olivia Kaplan, Sarah Schroeder, Bryan Hariadi, Simone Marrujo, Crina Curca Alec Salvino, Guillermo Gallareta Olivares, Ryan Koehler, Gary Geiss, Alexander Rosenberg, Charles Roco). These authors jointly supervised this work (Daniele Merico, Nima Alidoust, Hani Goodarzi, Johnny Yu). Corresponding authors: J.Y. H.G.

## Abstract

Building predictive models of the cell requires systematically mapping how perturbations reshape each cell’s state, function, and behavior. Here, we present *Tahoe-100M*, a giga-scale single-cell atlas of 100 million transcriptomic profiles measuring how each of 1,100 small-molecule perturbations impact cells across 50 cancer cell lines. Our high-throughput Mosaic platform, composed of a highly diverse and optimally balanced “cell village”, reduces batch effects and enables parallel profiling of thousands of conditions at single-cell resolution at an unprecedented scale.

As the largest single-cell dataset to date, *Tahoe-100M* enables artificial-intelligence (AI)-driven models to learn context-dependent functions, capturing fundamental principles of gene regulation and network dynamics. Although we leverage cancer models and pharmacological compounds to create this resource, *Tahoe-100M* is fundamentally designed as a broadly applicable perturbation atlas and supports deeper insights into cell biology across multiple tissues and contexts. By publicly releasing this atlas, we aim to accelerate the creation and development of robust AI frameworks for systems biology, ultimately improving our ability to predict and manipulate cellular behaviors across a wide range of applications.

## Introduction

A longstanding goal in cell biology is to build predictive, mechanistic models of how cells integrate signals and execute precise transcriptional and phenotypic responses (Endy and Brent 2001). Building expressive *in silico* models of cell behavior requires the generation of large and quantitative datasets that systematically map how cellular state (e.g., measured by its transcriptomic profile) is reshaped by a wide variety of interventions—including genetic, chemical, or environmental perturbations. Perturbative measurements elucidate causal gene-gene interactions, uncover feedback circuits, and expose compensatory pathways, thereby revealing the underlying regulatory networks governing cellular behavior and function (Pe’er 2005; Dixit et al. 2016; Adamson et al. 2016). Perturbative datasets built on modern high-throughput profiling techniques can be incredibly powerful for revealing basic biological processes with an unprecedented resolution, in turn enabling efforts to build *in silico* models of the cell (Bunne et al. 2024).

Traditionally, bulk transcriptomic measurements of cellular responses to perturbations have provided valuable insights (Bock et al. 2022; Jinek et al. 2012; Shalem, Sanjana, and Zhang 2015), but they often mask important heterogeneity within cell populations. Such heterogeneity may arise from differences in cell cycle phase, baseline transcriptional programs, or genetic background, leading to diverse cellular outcomes even under identical external conditions (Buettner et al. 2015; Trapnell et al. 2014; Macaulay et al. 2016). Single-cell assays – such as single-cell RNA sequencing (scRNA-seq) – have revolutionized our ability to resolve this heterogeneity, enabling the molecular characterization of individual cells with unprecedented detail (Macosko et al. 2015; Aldridge and Teichmann 2020).

Studies leveraging scRNA-seq have expanded our understanding of how cells respond to a wide array of perturbations, ranging from small-molecule treatments (Srivatsan et al. 2020) to pooled CRISPR-based genetic interventions (Replogle et al. 2022). Complementary advances in combinatorial scRNA-seq (Rosenberg et al. 2018; Xie et al. 2024; Cao et al. 2017; Datlinger et al. 2021) and next-generation sequencing (Simmons et al. 2023; Almogy et al. 2022) further enhance throughput and cost-efficiency, pushing feasibility of multi-million cell datasets.

Despite these advances, the availability of large-scale, perturbation-focused single-cell datasets remains limited. Although some atlas-scale efforts have begun to aggregate data from multiple studies, challenges such as batch effects, inconsistent protocols, and limited coverage of perturbations persist (Z. Li et al. 2024; Tyler et al. 2024). The low number of observations for a given cell-perturbation combination also restricts the power to detect robust patterns, particularly for subpopulations of interest (Tyler et al. 2024; Chen et al. 2022). Furthermore, a significant fraction of existing data focuses on a narrow set of perturbations — often aligned with specific disease or drug discovery goals — leaving many questions about the broader perturbation landscape unanswered.

To address these challenges and enable next-generation computational modeling of cellular function, we developed the Mosaic platform (J. X. Yu et al. 2024). Mosaic exploits single-cell RNA sequencing (scRNA-seq) to capture high-resolution transcriptomic responses to thousands of perturbations in parallel. By constructing “cell villages” (Mitchell et al. 2020; C. Yu et al. 2016; McFarland et al. 2020) of tens to hundreds of distinct cell models in each experiment followed by genetic demultiplexing, the platform substantially reduces batch effects while scaling to an increasingly larger number of perturbation sets.

Leveraging Mosaic, we generated *Tahoe-100M*, the largest publicly available single-cell dataset to date. *Tahoe-100M* comprises over 100 million single-cell transcriptomes collected from 50 cancer cell lines subjected to more than 1,100 small-molecule perturbations at varied doses. Though we apply pharmacological agents here, the underlying framework is broadly applicable to diverse perturbations, facilitating both fundamental investigations of cell biology and more translational applications such as drug discovery. By publicly releasing *Tahoe-100M*, we aim to catalyze the development of robust, context-aware AI models of cellular behavior, ultimately empowering the scientific community to gain deeper insights into gene function, regulatory networks, and their modulation by environmental cues.

## Results

### Generating the *Tahoe-100M* Atlas using Tahoe Therapeutics’ Mosaic platform

We leveraged our Mosaic platform to measure the transcriptomic responses to drug treatments across a panel of 50 established and commercially-available cancer cell lines (**Figure 1A**). For this, we cultured spheroids consisting of a mixture of cell lines in suspension and subjected each spheroid to individual treatments, including DMSO vehicle controls. After 24 hours of drug treatment, spheroids were dissociated, fixed, and profiled using the Parse GigaLab kit, a scalable scRNA-seq assay enabled by combinatorial barcoding (Rosenberg et al. 2018). In addition to generating single-cell gene expression matrices across treatments, we detected genetic variants present in scRNA-seq libraries and performed SNP-based deconvolution to assign cells to their cell line of origin (**Figure 1B**) (Kang et al. 2018).

**Figure 1.**
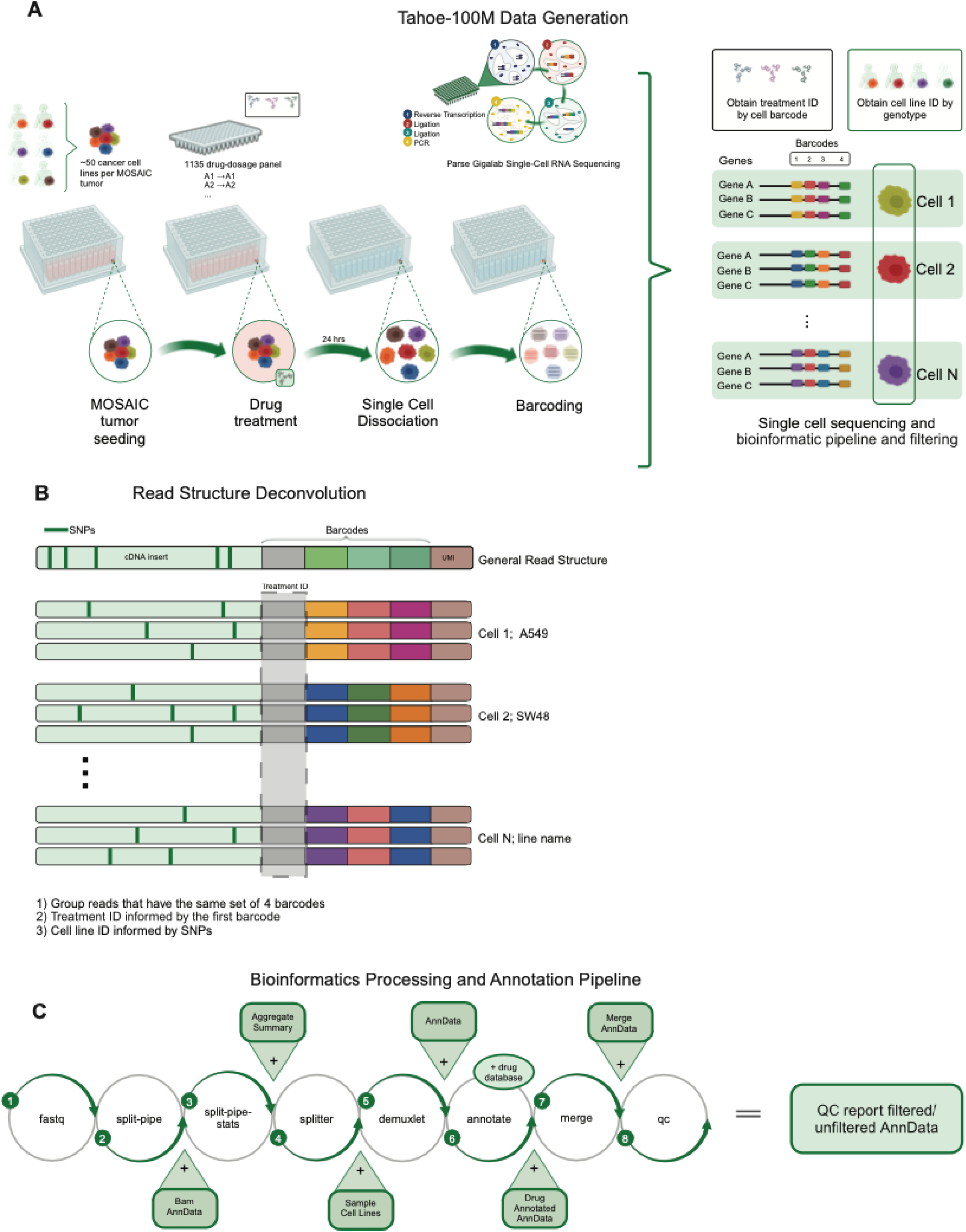
Mosaic platform: the experimental and informatics workflow used to generate *Tahoe-100M*. **(A)** Experimental schematic: each Mosaic tumor is composed of 50 cancer cell line models, which are seeded into 96-well plates. Each well receives a drug perturbation, and after 24h, tumors are dissociated into single cells and barcoded using the Parse GigaLab kit. These barcodes, along with known genotypes of each cell line, then allow for treatment and cell line deconvolution. **(B)** Read structure and bioinformatic pipeline filtering. Each read consists of a cDNA insert, four barcode regions, and a unique molecular index (UMI). After grouping together reads by those that share the same four barcode regions, drug perturbation identity can be resolved by the first barcode region, and cancer model identity resolved by the SNPs present in the cDNA insert. **(C)** Deconvolution and bioinformatics pipeline to process, annotate, and QC cells, resulting in a single-cell gene expression matrix with assigned treatment and cell line identity.

For this dataset, a total of 14 96-well plates of spheroids were seeded and drug treated. During library preparation, barcoded single cells from each plate were pooled and distributed into equal cell count “sublibraries” such that each sublibrary contains representation from all samples on a given plate but has unique sequencing adapters.We processed a range of between 72 to 165 sublibraries per 96-well plate of spheroids, resulting in a total of 1,786 sublibraries (**Table 1**). After sequencing, we obtained ∼1.4 trillion raw sequencing reads representing a total of 153 million cells (between 8.2 and 14.5 million cells from each plate). Cells had a mean of 2,288 transcripts (median of 1,890) with some variation at per-sublibrary level based on sequencing depth. After processing with Tahoe therapeutics’ filtering criteria (Methods), we recovered 100.6 million cells that pass minimal filters and 95.6 million cells that pass full filters (**Methods**; **Figure 1C**). We used the latter set for all the downstream analyses presented here.

**Table 1.** Per-plate breakdown of key statistics in *Tahoe-100M*.

| Plate | Number of Sublibraries | Total cells | Cells passing full filters | Median transcripts/cell | Mean transcripts/cell |
| --- | --- | --- | --- | --- | --- |
| 1 | 72 | 8228141 | 5299930 | 1953 | 2386 |
| 2 | 126 | 11645073 | 7687184 | 2098 | 2563 |
| 3 | 105 | 8653781 | 4158278 | 1462 | 1910 |
| 4 | 112 | 10570863 | 6679884 | 1918 | 2300 |
| 5 | 144 | 11280770 | 5908471 | 1647 | 2097 |
| 6 | 154 | 11032563 | 7285856 | 2159 | 2779 |
| 7 | 130 | 9916566 | 5227768 | 1571 | 1978 |
| 8 | 158 | 12564738 | 8516367 | 1929 | 2375 |
| 9 | 111 | 9082806 | 5569622 | 1727 | 2174 |
| 10 | 142 | 11878188 | 7731986 | 1841 | 2237 |
| 11 | 110 | 11352556 | 7062820 | 1761 | 2100 |
| 12 | 156 | 14515762 | 10188591 | 2019 | 2513 |
| 13 | 165 | 12746471 | 8106500 | 1863 | 2267 |
| 14 | 101 | 9446911 | 6201077 | 2025 | 2357 |

### Cell line and treatment diversity in *Tahoe-100M*

Of the 50 cell lines in *Tahoe-100M*, 47 had sufficient representation across the conditions to be evaluated in downstream analyses. These 47 lines are derived from 13 different organs (majority from lung, bowel, pancreas and skin) and carry a diverse set of driver mutations, with TP53, KRAS and CDKN2A altered in about half of the cell lines (**Figure 2A**). Across all 379 distinct drugs used in the atlas, 180 are classified into 25 mechanisms of action (MOA), with a median of 5 unique drugs per MOA (maximum 27, minimum 3) (**Figure 2B-C**). The majority (69%) of these drugs are approved agents and the compound classes target a large diversity of cancer-relevant pathways. These drugs reportedly target 325 genes (**Methods)**, with 120 genes being targeted by more than one drug (**Figure 2C**). This dataset captures 17,813 unique cell line-drug conditions, representing a 31-fold increase in the number of drug-perturbed conditions compared to a benchmark single-cell drug screen (Srivatsan et al. 2020) and a 29-fold increase in the number of observations per condition compared to a benchmark genetic perturbation screen (Replogle et al. 2022) (**Figure 2D**).

**Figure 2.**
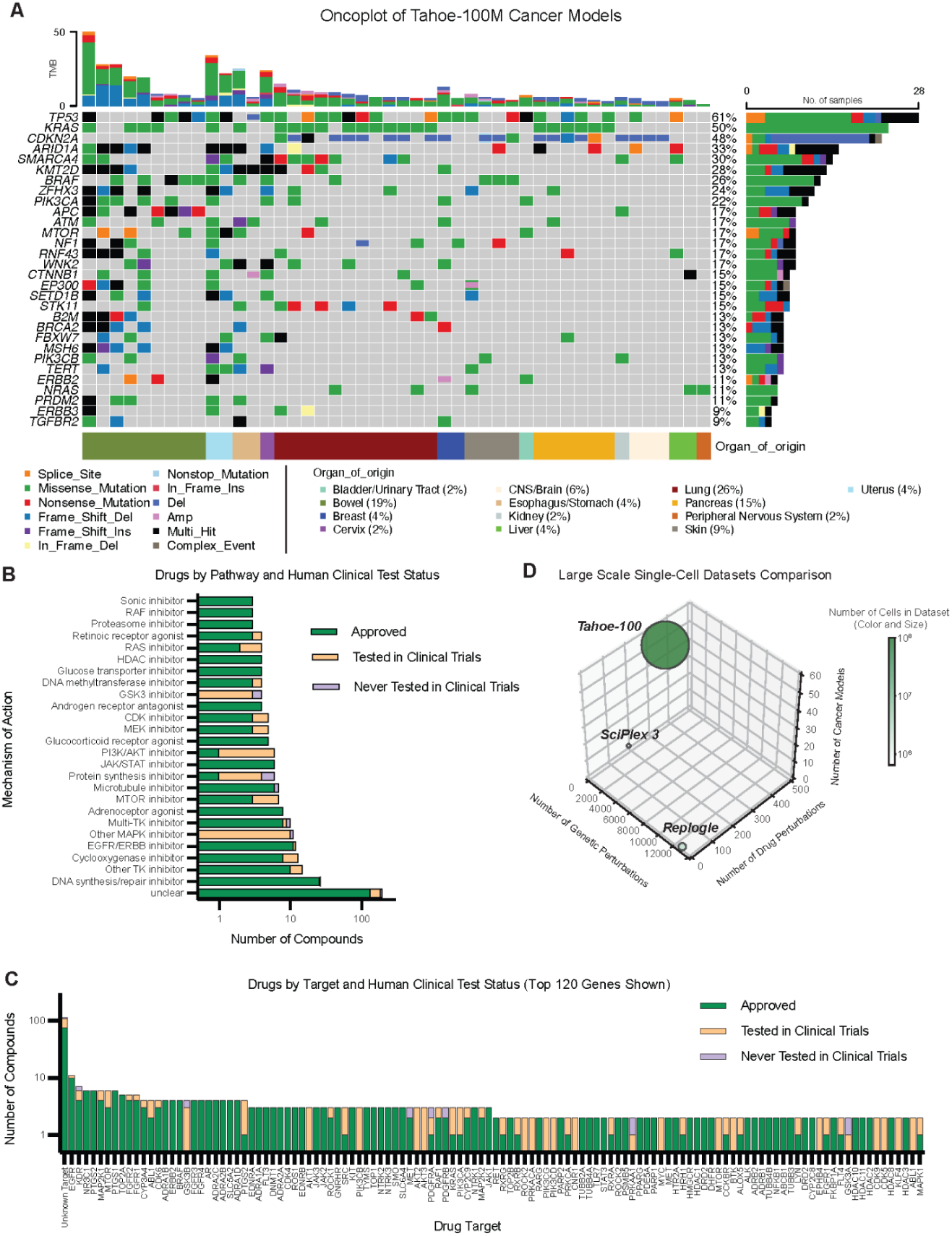
Diversity of cellular models and drug perturbations in *Tahoe-100M*. **(A)** Oncoplot displaying the distribution of the top 30 driver mutations found amongst the top 47 cellular models by abundance in *Tahoe-100M*. The mutations are further classified by type and the cellular models are further classified by organ of origin. **(B)** Mechanism of action grouping of the 379 drug perturbations in *Tahoe-100M*, further stratified by clinical trial status. **(C)** Target gene grouping of the 379 drug perturbations in *Tahoe-100M*, further stratified by clinical trial status, shown for the top 120 out of 325 target genes. **(D)** Comparison of number of perturbations, unique cellular models, and total cell count to other benchmark perturbational single-cell RNA-sequencing datasets.

### The global transcriptomic landscape captured by *Tahoe-100M*

We began our analysis of *Tahoe-*100M by performing dimensionality reduction with scVI (Lopez et al., 2018; Gayoso et al, 2022) to learn a 10-dimensional embedding for each cell. Given the challenge in visualizing a dataset of 100M cells, we sampled 140,000 high quality cells from each of 47 cell lines (6.58 million cells total), computing 2-D tSNE coordinates, and plotting a random selection of 200,000 cells (**Figure 3A-C**). Also evident in the tSNE visualization, we noted a clear separation of cells based on their genetic identity (and cell cycle phase) rather than their plates of origin in the transcriptomic space, suggesting a lack of substantive batch effects across this unified ∼100M cell atlas (**Figure 3A-C**). Regardless, the differential gene expression analyses described in the following sections account for the possible batch effect by leveraging the DMSO-treated wells included in all plates.

**Figure 3.**
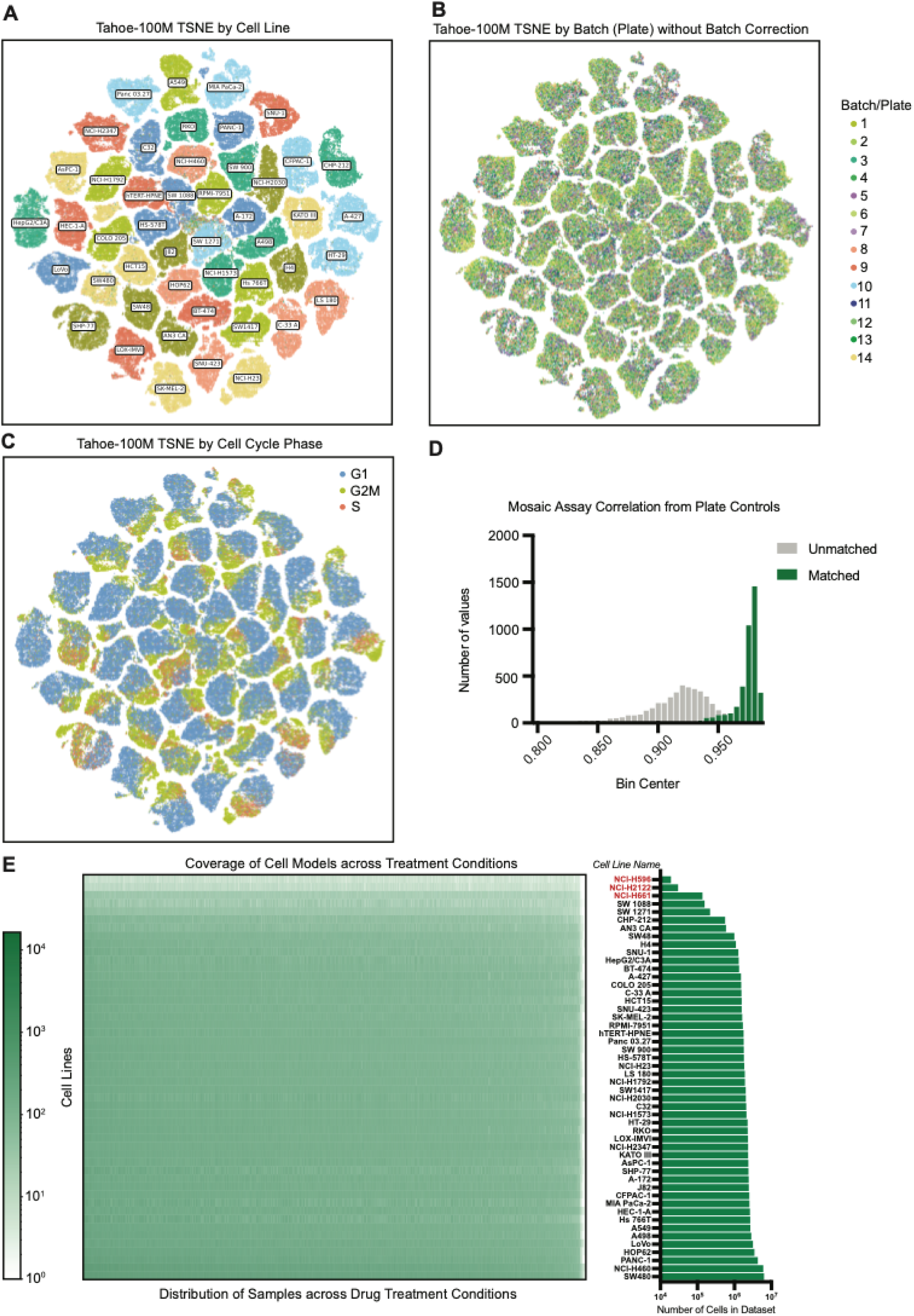
Visualization of absolute gene expression in *Tahoe-100M*. **(A)** 2D-tSNE representation of the SCVI model, plotting a random selection of 200,000 cells with equal contribution from the top 47 cell lines by abundance. Clusters are annotated by cancer model identity. **(B)** Same as in (A) but cells annotated by the 96-well plate of Mosaic tumors of origin. **(C)** Same as in (A) but cells annotated by their cell cycle phase. **(D)** Distribution of Pearson correlation values for pairwise gene expression vector correlations between the two biological replicate 96-well plates in the Atlas. Cells were aggregated into pseudobulk gene expression vectors based on matching cancer model identity and drug perturbation. 4000 unmatched and matched cancer model identity and drug perturbation comparisons were randomly selected. **(E)** Left: Heatmap of cell coverage across all Mosaic tumors in the Atlas, colored by number of single-cell observations for a given cell line and drug-dosage treatment. Right: Total single-cell counts for all 50 cancer models in the *Tahoe-100M*, with the 3 lowest abundance cell lines highlighted in red.

Taken together, after applying our QC filtering, we obtained a dataset comprising 47 cell lines, 379 drugs, 1,135 drug-dose combinations and 52,886 unique cell line-drug-dose conditions, with a median of 1,287 cells per condition. To highlight the reproducibility of the Mosaic platform, we included Plate 14 as a biological replicate of Plate 6. Using the untransformed gene expression matrices from these plates, we performed a pseudobulk grouping of all cells belonging to the same plate, cell line, and drug concentration treatment and calculated the Pearson correlation across these pseudobulk gene expression vectors. Reassuringly, we observed higher correlation between conditions with matched treatment and cell identity (estimate q25: 0.97, median: 0.975, q75: 0.98) compared to unmatched comparisons between the two plates (estimate q25: 0.892, median: 0.915, q75: 0.929) (**Figure 3D**).

### Drug-induced transcriptomic effects across cell lines and mechanisms of action

To quantify technical and biological factors contributing to cell groupings in *Tahoe-100M*, we estimated the Local inverse Simpson’s index (LISI) based on different metadata factors over a subsample of the data. By repeating this process across 100 different subsamples, we quantified the mean and standard deviation of the estimates of median LISI (**Figure 4A**). Consistent with our expectations and the tSNE visualizations, these results suggest that cell line identity, followed by cell cycle phase, are the most major factors in driving stratification of the data. And as expected, drug treatment and dose have a more modest effect relative to these factors, suggesting that differential gene expression analysis is required to better delineate drug-induced changes.

**Figure 4.**
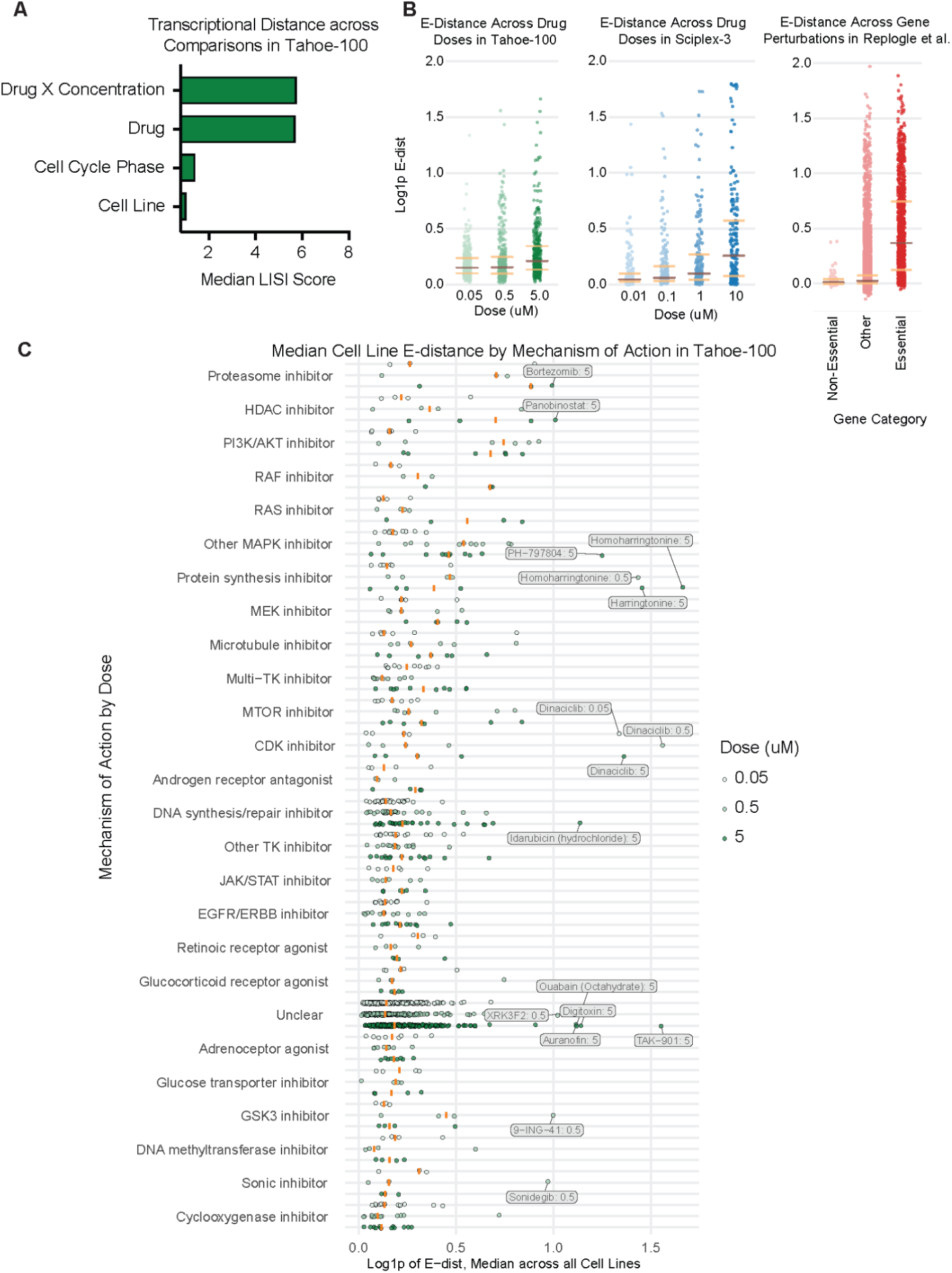
Overall impact of treatments on the transcriptome across cell lines in *Tahoe-100M*. **(A)** Local inverse Simpson’s index (LISI) scoring based on several selected categorical metadata variables. **(B)** Comparison of median log1p E-distance between perturbed and control cells of the same cellular identity in the *Tahoe-100M* and two other benchmark single-cell perturbational Atlas datasets. The median E-distance across all cellular models was plotted. Left: *Tahoe-100M*, stratified by drug dosage. Middle: SciPlex-3 dataset, stratified by drug dosage. Right: Replogle et al. Perturb-seq dataset, stratified by classification of perturbed gene. **(C)** Comparison of median log1p E-distance between perturbed and control cells of the same cellular identity in *Tahoe-100M*, taking the median value across all cellular models and grouping by drug mechanism of action (MOA). MOAs were sorted by high to low median value at the highest dose.

To explore the landscape of transcriptomic responses to drug treatments, we first investigated the magnitude of effect for a given treatment across the entire transcriptome using E-distance (Peidli et al. 2024). This distance metric, which is well-established in single-cell perturbation data analysis, quantifies the separability of a perturbed population from its controls (Norman et al. 2019, (Replogle et al. 2022)). We calculated E-distances for all drug conditions within a cell line using our scVI embeddings (**Methods**) and consistent with our expectations, we observed a larger median E-distance at the highest drug doses across treatments (**Figure 4B**, left). A similar E-distance distribution was observed when applying this analysis to the previously published single-cell perturbation dataset Sciplex3 (Srivatsan et al. 2020) (**Figure 4B**, center). Moreover, when examining genome-wide CRISPRi perturbations (Replogle et al. 2022), and stratifying genes by essentiality, we found drug perturbations across this dataset to exhibit an intermediate effect (**Figure 4B**, right).

We observed several notable outliers when stratifying drugs by their reported MOA (**Figure 4C**). For instance, harringtonine and homoharringtonine are plant-derived alkaloids that induce cancer cell cytotoxicity by directly inhibiting ribosome polypeptide elongation (Kantarjian et al. 2001), as opposed to modulating translation factors or upstream signaling pathways like other protein synthesis inhibitors in this study. Additionally, dinaciclib is a potent and selective inhibitor of CDK1, CDK2, CDK5 and CDK9, which inhibits cell proliferation in a broad set of cancer models (Parry et al. 2010). Aside from these outlier drugs, proteasome inhibitors, HDAC inhibitors and PI3K/AKT inhibitors tended to have larger effects across dose levels. More generally, anti-cancer drugs active in epithelial solid tumors (MAPK, PI3K/AKT, MTOR and CDK inhibitors, microtubule-interfering agents and DNA synthesis/repair inhibitors) tended to have larger E-distances than other drugs (e.g. retinoic receptor agonists, adrenoceptor agonists, cyclooxygenase inhibitors) or unclassified MOA.

For dimensionality reduction based on differential expression gene-set scores (**Figure 5A**), we opted to consider each unique combination of cell line, drug and dose as a single data point. The presence of drugs with large E-distances were reflected in the 2-dimensional t-SNE and nnMDS, with both showing a clear E-distance gradient from the center to the periphery (**Figure 5B-C**). Considering MOAs with larger E-distance and more relevant to epithelial cancers (HDAC inhibitors, proteasome inhibitors, PI3K/AKT inhibitors, etc), we observed a high degree of overlap yet some separation, especially in regions with higher E-distance (**Figure 5D-E**). Finally, we focused on the KRAS G12C-specific covalent inhibitor Adagrasib and the more mutation agnostic Pan-RAS inhibitor RMC-6236, showing a clear separation of RMC-6236 and Adagrasib in KRAS G12C cell lines relative to Adagrasib in non-KRAS G12C cell lines, following expectations (**Figure 5F-G**).

**Figure 5.**
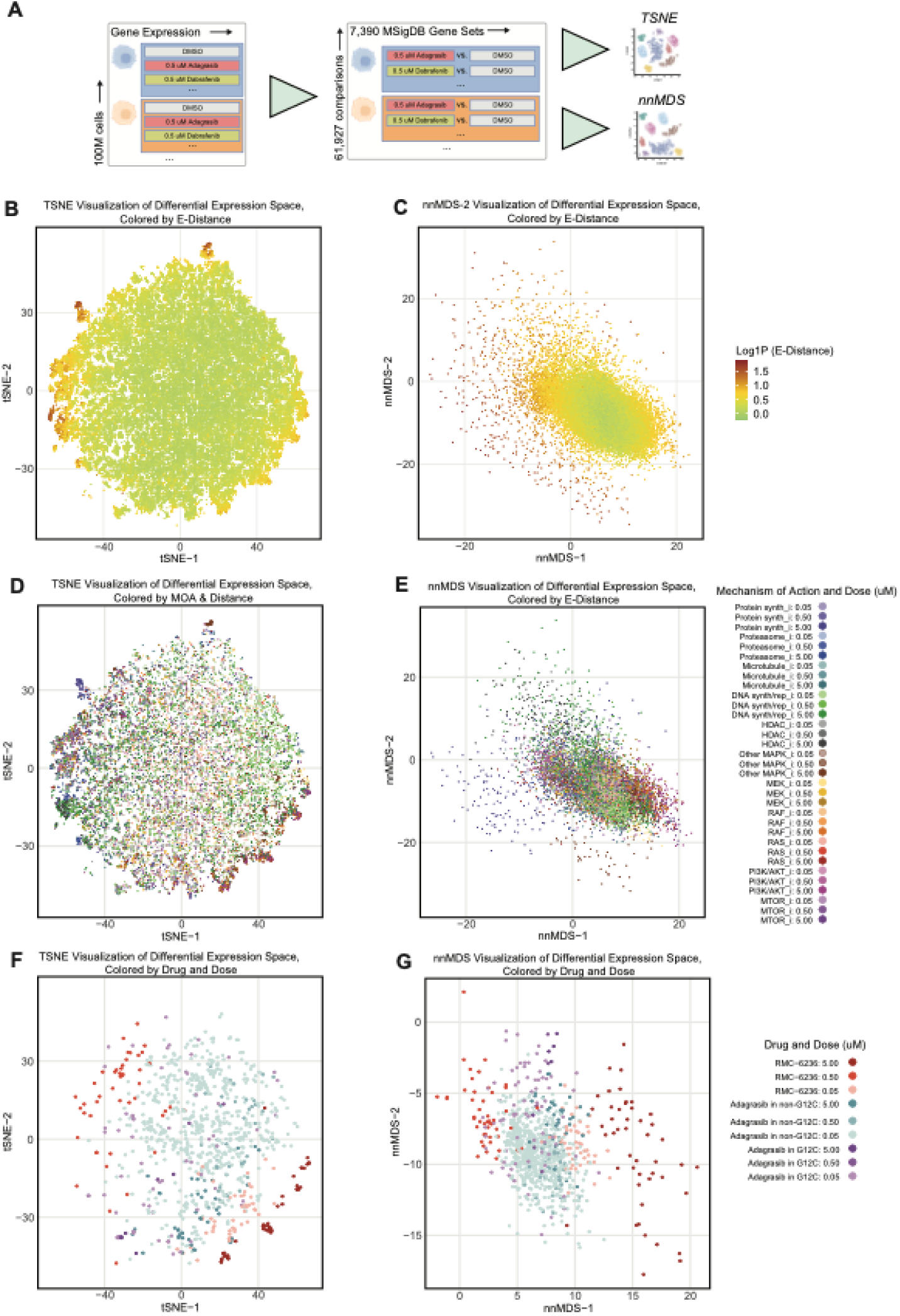
Differential gene set expression scores visualized using dimensionality reduction of pseudobulk cellular models and drug treatment conditions. **(A)** Single cell datapoints in the Atlas were grouped in pseudobulk by shared cellular model and drug treatment conditions, from which differential gene expression scores were calculated for 7,390 gene sets. The resulting matrix of 61,927 cellular model and drug treatment pseudobulk groups and 7,390 gene sets was used as input to two different dimensionality reduction visualizations. **(B)** 2D tSNE visualization of the matrix in (A) colored by E-distance. **(C)** 2D nnMDS visualization of the matrix in (A) colored by E-distance. **(D)** Same tSNE as in (B) colored by drug mechanism of action (MOA) and dosage and removing points belonging to drugs for which there was no available MOA annotation were filtered from the plot. **(E)** Same 2D nnMDS visualization as in (C) colored by drug MOA and dosage, restricting to the MOAs with largest e-distance. **(F)** Same 2D tSNE as in (B), only visualizing points that correspond to RAS inhibitors RMC-6236 and Adagrasib, colored by dosage and, for Adagrasib only, by presence/absence of a KRAS-G12C mutation in the cell line. **(G)** Same 2D nnMDS as in (C), same coloring convention as (F).

### Context-dependent inhibition of the RAS/RAF pathway

To further investigate the transcriptional effects of drugs, we focused on targeted treatments that modulate the RAS/RAF pathway. We used *Vision* (DeTomaso et al. 2019) to compute gene-set expression scores at the cell level and then computed a differential score between aggregated treatment and control conditions. We adopted a MSigDB.c6 gene-set (Subramanian et al. 2005) that we previously identified as a highly effective transcriptional signature for RAS/RAF pathway activity. We inspected differential *Vision* scores (Methods) for this signature for the BRAF-V600E inhibitor Dabrafenib across three cell line stratifications: those with KRAS mutations in the absence of additional RAF or RAS mutations; those with BRAF-V600E mutations lacking concurrent RAS or RAF mutations; and those devoid of both RAF and RAS mutations. Notably, treatment with Dabrafenib elicited the most pronounced effect in cell lines harboring BRAF mutations, as evidenced by significantly lower scores at both the lowest (P = 0.0013, Welch’s t-test) and highest (P = 0.00023, Welch’s t-test) doses, whereas KRAS-mutated lines showed minimal change (**Figure 6A**). Given that KRAS activating mutations occur upstream of BRAF activating mutations, these findings align with the expected dynamics of the RAS/RAF signaling pathway. In contrast, treatment with RMC-6236, a pan-RAS inhibitor, produced the greatest effect in the KRAS-mutated group, with the most substantial difference in Vision scores observed at the lowest (p = 0.065, Welch’s t-test) and intermediate (p = 0.063, Welch’s t-test) doses compared to the unmutated RAS group (**Figure 6B**). These results underscore the KRAS-specific action of RMC-6236, which reduces KRAS signaling in KRAS-mutated cell lines.

**Figure 6.**
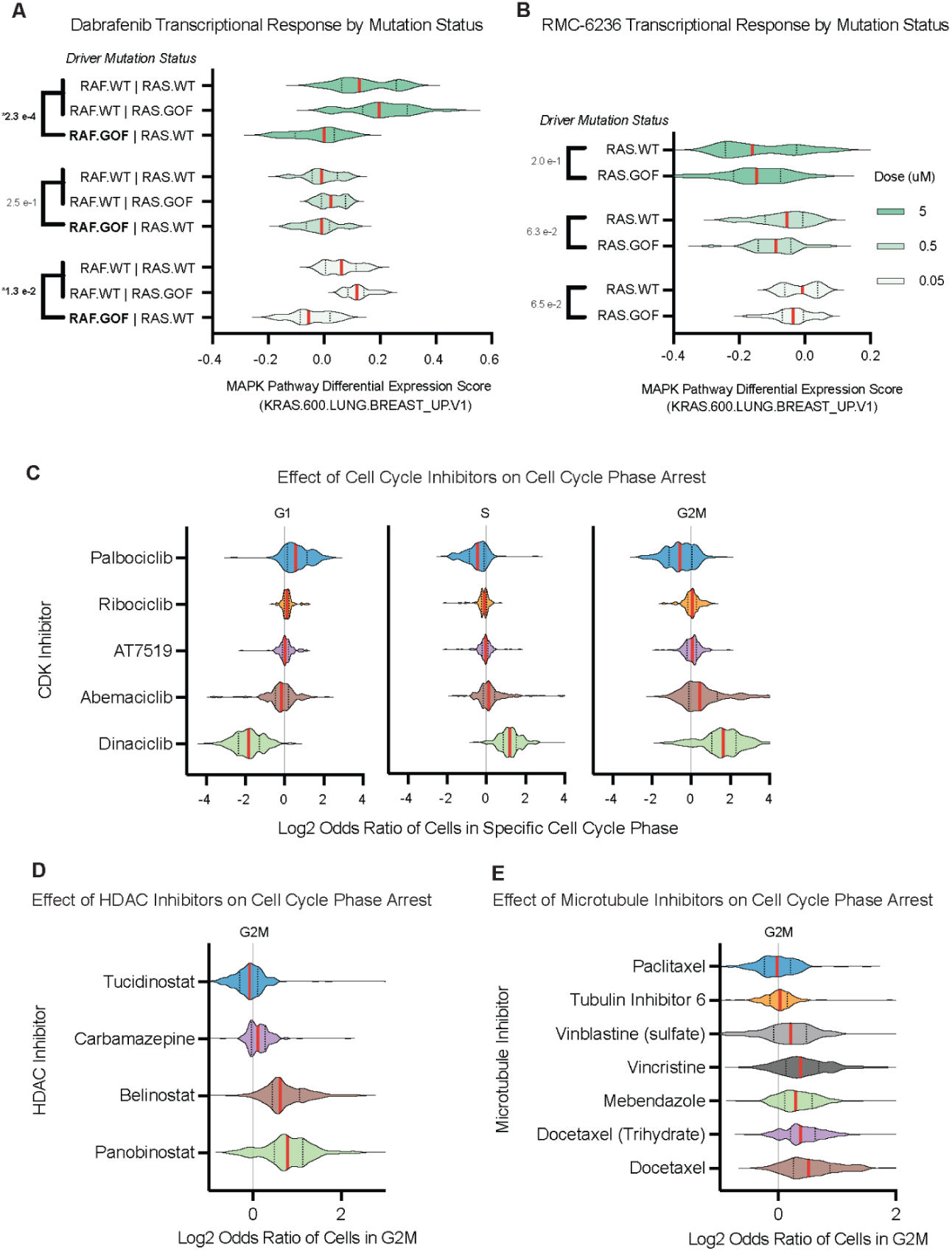
Gene expression and cell cycle phase specific responses to selected drugs. **(A)** Violin plot of differential gene expression for a 600-gene transcriptional signature of RAS/RAF pathway activity for Dabrafenib, a RAF inhibitor, grouped by cellular model genotype (wild type or gain of function) and by drug dosage (5.0, 0.5, 0.05 uM top to bottom). **(B)** Violin plot of differential gene expression for a 600-gene set implicated in the MAPK pathway for RMC-6236, a RAS inhibitor, grouped by RAS mutational status and by drug dosage (5.0, 0.5, 0.05 uM top to bottom). **(C)** Comparison using log_2_-odds ratio of each of the cell cycle phase proportions across all observations from a given cellular model in response to a selected set of cell cycle inhibitors as compared with the untreated control. Each of 47 cellular models and all 3 drug concentrations were plotted on the same violin plot. **(D)** Comparison of log_2_-odds ratio of cells in G2M phase for selected HDAC inhibitors. **(E)** Comparison of log_2_-odds ratio of cells in G2M phase for selected microtubule inhibitors.

We subsequently investigated the impact of drug treatments on cell cycle phase distribution by computing the odds ratio for the percentage of cells in each phase relative to the control condition. Within the class of CDK inhibitors, we observed that Palbociclib and Dinaciclib exhibited distinct enrichments, with Palbociclib showing enrichment in the G1 phase and Dinaciclib in the G2/M phase (**Figure 6C**). This is in contrast to the other CDK inhibitors — Abemaciclib, AT5719, and Ribociclib — which displayed little to no enrichment. Notably, Dinaciclib inhibits CDK1, CDK2, CDK5, and CDK9 and has been shown to induce G2/M phase arrest, whereas Palbociclib selectively targets CDK4/6 — responsible for phosphorylating the retinoblastoma protein (Rb) and facilitating the transition from G1 to S phase (Rajput et al. 2016; Crozier et al. 2022). Further investigation of the G2/M phase enrichment revealed that HDAC and microtubule inhibitors induced the greatest G2/M arrest. Within the HDAC inhibitors, Belinostat and Panobinostat predominantly drove this G2/M enrichment, in contrast to Carbamazepine—which, despite its role as a histone deacetylase modulator, showed little enrichment—and Tucidinostat (Chidamide), which demonstrated a negative enrichment (**Figure 6D**).

The absence of enrichment observed under Carbamazepine conditions may be attributed to its primary role as an anti-epileptic agent, which functions predominantly as a voltage-gated channel modulator and exhibits only modest HDAC inhibitory activity (Beutler et al. 2005). In contrast, the negative enrichment observed with Tucidinostat is likely due to its inhibition of HDAC1, HDAC2, HDAC3, and HDAC10, a mechanism that has been shown to induce G0/G1 cell cycle arrest through the downregulation of CDK4 (Beutler et al. 2005; Zhong et al. 2021). With respect to microtubule inhibitors, these agents are believed to induce mitotic arrest via activation of the mitotic checkpoint, thereby ensuring proper chromosomal alignment with the spindle apparatus prior to anaphase onset (Čermák et al. 2020). Overall, the majority of these agents produced a general enrichment in the G2/M phase, with the notable exceptions of Paclitaxel and Tubulin Inhibitor 6, which exhibited minimal enrichment (**Figure 6E**).

## Discussion

In this study, we present *Tahoe-100M*, a large-scale single-cell perturbation atlas that far surpasses previous datasets in both breadth and depth, encompassing over 100 million single-cell transcriptomes from 50 cell lines treated with more than 1,100 small-molecule agents. By capturing transcriptomic responses at single-cell resolution, *Tahoe-100M* reveals how diverse pharmacological perturbations, across multiple genetic backgrounds, reshape cellular states. The unique scale and design of this resource address key challenges in systems biology, particularly the need for comprehensive perturbation data at scale that can power next-generation AI-driven models of the cell (Bunne et al. 2024).

### Enabling AI-driven models of the cell with a large, single-cell perturbation atlas

1. *Diverse training data*: A broad and balanced set of conditions and drug MOAs ensures that AI models see complex, context-dependent gene regulatory modulations rather than uniform or highly specialized responses.
2. *Deep perturbation coverage*: Multiple drugs that converge on the same pathway or target allow models to learn finer-grained distinctions in mechanism of action, reinforcing a more robust representation of gene-gene and gene-drug interactions.
3. *Context-specific insights*: Perturbations performed in a multiplexed “cell village” framework unify the data across multiple cell lines and tissues, enabling AI systems to discern how genetic and epigenetic contexts modulate drug efficacy and cell-state transitions.

Together, these features provide a strong foundation for self-supervised, semi-supervised, and transfer learning approaches, in turn opening up new avenues for building *in silico* models of the cell. The success of AI frameworks in fields like natural language processing and image recognition highlights how performance can scale dramatically with increased data size and diversity. We anticipate a similar trajectory for single-cell biology as *Tahoe-100M* provides a critical leap in the availability of meaningful training data.

### Inter- and intra-population cellular heterogeneity in drug response

*Tahoe-100M* also exposes the profound heterogeneity in how cells respond to identical pharmacological perturbations, unveiling important considerations for cancer therapy:

1. *Subpopulation dynamics*: Single-cell resolution reveals that even within a genetically uniform cell line, multiple transcriptional substates can exist that each exhibit different susceptibility to

a given drug (Dann et al. 2022; Skinnider et al. 2021). We observed, for instance, distinct distributions of cell cycle state that varied across drug classes, underscoring the context-dependent interplay between baseline cellular programs and drug perturbations (Weinberger, Lin, and Lee 2023; Schiebinger et al. 2019; Dong et al. 2023; Papalexi et al. 2021).

1. *Pathway-specific effects*: Our results show that targeted inhibitors, such as those directed against RAS or RAF, elicit markedly different transcriptomic shifts in a driver

mutation-dependent manner. This suggests that single-cell profiles can help pinpoint which subsets of cells are poised to evade treatment via compensatory pathways, and can also identify instances in which pathway crosstalk complicates straightforward inhibitory strategies.

1. *Predictive biomarkers*: By encompassing a broad panel of cell lines, *Tahoe-100M* enables the systematic identification of transcriptional correlates linked to drug response. These correlates can serve as potential biomarkers, guiding patient stratification and helping clinicians tailor treatments to molecular profiles most likely to respond (C. Yu et al. 2016; Tsherniak et al. 2017).

Understanding these nuances of drug-cell interactions is central to dissecting how treatment resistance emerges and persists. Moreover, this heightened clarity about cellular heterogeneity helps refine AI models of drug response, which must account for the full distribution of potential outcomes rather than an “average” or bulk response.

### Accelerating cancer drug discovery and translational research

Cancer remains one of the most complex diseases to treat, given its high genetic diversity and the multiple layers of co-opted regulatory dynamics in tumor cells. Data generated by our Mosaic platform highlights the following opportunities for translational research:

1. *Context-aware drug development*: By capturing how each drug interacts with varied genetic backgrounds, *Tahoe-100M* offers an actionable blueprint for discovering novel therapeutics. Researchers can rapidly mine this atlas to identify compound classes that exploit specific mutational contexts or that can circumvent known resistance mechanisms.
2. *Combination therapies*: The atlas serves as a starting point to investigate synergy between different classes of drugs. Single-cell profiling of combination regimens could reveal whether targeting multiple vulnerabilities—e.g., simultaneous inhibition of CDK and HDAC pathways—yields more durable responses across heterogeneous cell populations.
3. *Insights for personalized medicine*: In an era moving toward precision oncology, linking genotype-specific transcriptional responses to high-throughput drug screens could guide personalized treatment regimens. Cell lines that mirror certain patient subtypes can be interrogated to discover or validate effective drug cocktails, informed by differential expression, cell cycle arrests, or pathway activation states.

### Future directions

By publicly releasing *Tahoe-100M*, we aim to seed a productive cycle of data-driven discovery and AI model development. Researchers can use this resource not only to improve existing computational frameworks but also to unlock novel insights into fundamental cell biology, drug efficacy, and therapeutic resistance. As the field coalesces around large-scale perturbative atlases, the synergy between experimental and computational innovations will move us closer to a predictive, mechanistic understanding of the cell—and ultimately, to more effective, personalized treatments for patients.

## Methods

### Cell culture

Cell lines were sourced from ATCC or NCI-43 and cultured in complete 1X RPMI-1640 medium (Cytiva) supplemented with 10% FBS and 1.5x pen-strep at 37C and 5% CO2. Cells were harvested using TrypLE Express (Gibco) if adherent or directly from the media if suspension-based and pooled together at equal ratios, combined with Cultrex (R&D Systems), and cast into 14 total deep-well 96 well plates (Diaago).

### Compound sourcing and treatment

Compounds were sourced from MedChemExpress and formulated by diluting the compound to final concentration in 1X RPMI-1640 complete, 10% FBS, 1.5x pen-strep with no more than 1% DMSO in the final solution. 14 96 well drug media plates were formulated, with well H11 and H12 on each plate consisting of control media with no compounds. Drug media was overlaid on 3D cell cultures, matching 96 well plate coordinate and plate number. Cells were exposed to drug media for 24 hours at 37C and 5% CO2.

### Single cell dissociation and fixation

3D cell cultures were dissociated into single cells using TrypLE Express in 96 well plates. Samples were then fixed following the High-Throughput Evercode Cell Fixation v3 kit (Parse Biosciences) and stored in 96 well plates. Throughout both processes, 96 well plate coordinate and plate number identity was maintained for samples, resulting in an output of 14 96 well plates containing fixed cell suspensions. Fixed samples were stored at −80C and subsequently shipped on dry ice to the Parse Biosciences GigaLab.

### Barcoding

Samples were diluted such that roughly the same number of cells from each sample were input into barcoding. Each plate of fixed cells underwent three rounds of combinatorial barcoding using Section 1 of the Parse Biosciences GigaLab protocol. After barcoding, cells were distributed into individual sublibraries of ∼100,000 cells and lysed according to Parse Biosciences GigaLab protocol.

### Sublibrary Preparation

Lysates were processed through Sections 2 and 3 of the Parse Biosciences GigaLab protocol in batches of 96 with the Hamilton Vantage liquid handling robot. At the end of Section 3, an Illumina i7 and i5 index was added to each sublibrary to produce a sequenceable molecule. Each sublibrary was then converted into an Ultima Genomics-compatible sequencing library as previously demonstrated (Simmons et al. 2023) and sequenced on the UG-100 sequencer.

### Sequencing

Sequencing was performed as described on Ultima Genomics sequencers (Almogy et al. 2022).

### Split-pipe processing

Fastq files generated from Ultima Trimmer workflow were processed through a modified version of spit-pipe v1.4.0 to accommodate libraries processed through the GigaLab. In addition to default options, we added the –yes_allwell option to aid in removing low quality cells from the final list of cell barcodes. For each sublibrary, split-pipe generated a cell by gene matrix (count_matrix.mtx) as well as relevant metadata for each cell (cell_metadata.csv) and a list of genes to which split-pipe maps fastq files (all_genes.csv). Gene and transcript definition was based on Ensembl release 109 and genome build 38 (https://ftp.ensembl.org/pub/release-109/gtf/homo_sapiens/Homo_sapiens.GRCh38.109.gtf.gz).

### Genotype-based cell line demultiplexing

Demultiplexing of the single-cell RNA-sequencing (scRNA-seq) data was conducted by assigning each cell to its respective cell line using demuxlet in conjunction with a curated reference set of single nucleotide polymorphisms (SNPs) derived across the cell lines (Kang et al. 2018). The reference set was constructed by filtering dbSNP to retain only exonic and untranslated region (UTR) variants with a gnomAD allele frequency greater than 0.1. These positions were subsequently processed with mpileup on the scRNA-seq data from each cell line to obtain genotype calls across all positions (H. Li 2011). The resulting SNP calls were then filtered to include only those with a depth (dp) greater than 6, mapping quality (MQ) exceeding 30, and quality score (Qual) above 50. These filtered variants were merged into a single VCF file, which was further refined by removing indels, multiallelic SNPs, and SNPs that were homozygous for the reference allele across all cell lines, yielding the final SNP reference. Benchmarking of this reference involved running demuxlet on a BAM file containing cells from known cell lines, serving as a ground truth. The performance evaluation demonstrated an accuracy exceeding 98%, with more than 90% of cells being labeled as singlets. It is worthwhile to note that demuxlet may designate cells as doublets when their profiles exhibit an equal likelihood of belonging to different cell lines; these classifications can represent either true doublets or cells with lower coverage or fewer SNPs.

### QC filters

Quality control (QC) was rigorously applied to all cells to ensure that only high-quality data were used for downstream analyses. QC filtering was implemented on two levels: full and minimal. The full filters required cells to have at least 700 unique molecular identifiers (UMIs), less than 20% mitochondrial reads, a UMI z-score within ±3 to remove outliers in total counts, a mitochondrial percentage z-score within ±3 to remove outliers in mitochondrial content, and at least 250 genes detected. In addition, only cells classified as singlets by demuxlet were retained, thereby excluding cells identified as doublets or ambiguous. The minimal filters applied were less stringent, requiring only a demuxlet call of singlet and less than 20% mitochondrial reads. For downstream analyses only cells passing full filters were used.

Further QC was performed at the condition level, where each cell line-drug treatment condition was required to have a minimum of 50 cells for inclusion in downstream analyses. Moreover, three cell lines (NCI-H661, NCI-H596, NCI-H2122) that consistently exhibited low counts across conditions were excluded from subsequent analyses.

### Replication analysis

UMI counts for single cells in plate 6 (7,285,856 cells) and plate 14 (6,201,077 cells) were aggregated into ‘pseudobulk’ UMI counts for each (cell line, treatment) pair (Squair et al. 2021), yielding 4,800 pseudobulk profiles for plate 6 and 4,796 for plate 14. The dataset was then restricted to the 4,796 (cell line, treatment) pairs common to both plates. The resulting pseudobulk UMI count matrices were concatenated, forming a dataset of 9,592 pairs.

Pearson correlation coefficients were computed for all pairwise comparisons of (cell line, treatment) pairs within plates, using log-transformed UMI counts (log(UMI counts + 1)) from the pseudobulk profiles.

Correlations were classified into two categories: with or without matched treatment and cell identity. 4,000 pairwise comparisons were randomly sampled from each category. These sampled comparisons were used to calculate the quartiles (25th, 50th, and 75th percentiles) of the Pearson correlation values for each category.

### Drug targets and mechanism of action (MOA)

We used MedChemExpress (https://www.medchemexpress.com/) and GPT-4o to annotate our list of compounds with known gene targets and mechanisms of actions (MOAs). We extracted compound descriptions and lists of targets (if known) as strings from MedChemExpress using Selenium.

To obtain targets, we first asked GPT-4o to propose a list of gene targets for each compound, given the compound name. We then asked the model to refine its proposed targets using the information from MedChemExpress, only keeping genes that were corroborated by the MedChemExpress information and translating gene names to standard symbols. If there were no corroborated targets, we asked the model to return the string “none”. Once we had a list of targets for each compound, we asked GPT-4o to consider the target list and the MedChemExpress information and determine if the compound acted on specific mutation(s) of any targets, returning that information separately.

We annotated MOAs in both “broad” and “fine” terms. To broadly annotate compounds as “inhibitor/antagonist” or “activator/agonist”, we asked GPT-4o to consider the MedChemExpress information and the target list for each compound and determine which category was appropriate, with the instruction to return “unclear” if there was not enough information to make the determination. To make finer MOA annotations, we provided GPT-4o with the same information, along with a manually-curated list of 25 mechanisms of action with some example compounds in each category. We asked the model to determine which of the MOAs was most appropriate for each compound, again with the instruction to return “unclear” if there was insufficient information.

We manually inspected the annotations for 97 of the 380 compounds. We noticed that some of the targets were not provided as standard gene symbols, so we ran the target list through GPT-4o again, asking it to correct this issue where present.

We also used GPT-4o to annotate whether these compounds were approved for human use and/or have been used in clinical trials. We prompted the model with the compound name and asked it to answer yes or no for each of the two questions, with an additional “notes” section in the answer dedicated to providing any important information for contextualizing the responses (e.g. if a specific enantiomer has not been approved for human use, but is used as part of a racemic mixture). We further iterated on the clinical trials annotations by asking the model to use clinicaltrials.gov to help corroborate its responses.

### scVI model

We set up an SCVI model (Lopez et al. 2018) to represent the expression levels of all 62,710 genes using 10-dimensional representations through a negative binomial likelihood with scvi-tools (Gayoso et al. 2022). To train the model, we collected 1,685 .h5ad AnnData files (Virshup et al. 2021), corresponding to individual ‘sublibraries’ from plates 1 through 13, into a single AnnCollection object with pointers to the AnnData objects loaded in ‘backed’ mode. Plate 14, a duplicate of plate 6, was excluded from training and reserved for validation and model criticism.

Cells passing full filters (95,624,334 cells) from plates 1 through 13 (89,423,257 cells) were included in training. These cells were further randomly split into 90% for training and 10% for validation. The model was trained for 10 epochs using single-cell observations, providing exposure to over 800 million examples of single-cell gene expression profiles.

Following training, the model and a corresponding AnnData file containing all cells passing full filters (including plate 14) were minified to enable interactive analysis (Ergen et al. 2024). In its minified format, the model provides gene expression estimates for 62,710 genes across 95,624,334 cells, utilizing 41 GB of storage.

### Single cell gene expression dimensionality reduction and visualization

Local inverse Simpson’s index (LISI) was used to quantify the spread or separability of SCVI cell transcriptome representations based on known factors (Korsunsky et al. 2019). LISI values were estimated using 30 nearest neighbors by randomly subsampling 10,000 cells, calculating the median LISI for each categorical factor, and repeating this process 100 times to obtain the mean and standard deviation of the median LISI estimates.

To visualize the gene expression data, three low-abundance cell lines were removed, and 140,000 cells per remaining cell line were uniformly sampled, yielding a representative subset of 6,580,000 cells with equal proportions per cell line. The posterior means of the SCVI representations for these cells were then visualized in two dimensions using the TSNE function from the cuML package in NVIDIA RAPIDS, with default parameters.

### Differential gene-set score calculations for dimensionality reduction

Genes were filtered to the intersection of genes in the reference genome and those present in gene set collections h.all, c5.GO, c6.all, and c2.cp from MSigDB (version v2023.1) (Subramanian et al. 2005), resulting in 20,250 genes. Standard deviations of normalized expression levels for each gene across the dataset were estimated by generating 100,000 samples of gene expression levels using the SCVI model and calculating the standard deviation of the sample per gene. Genes in the top 50% standard deviation were classified as ‘highly variable,’ totaling 10,126 genes.

The MSigDB gene sets were filtered to retain those containing between 15 and 500 genes (prior to any gene filter) and further refined to include only gene sets with at least 10 genes in the ‘highly variable’ set, resulting in 7,390 retained gene sets. Gene set activity was quantified using an adapted version of Vision scores (DeTomaso et al. 2019) applied to normalized expression levels from the SCVI model. The Vision score for a gene set was computed as the logarithm of the geometric mean of SCVI-normalized gene expression levels within the set.

To assess drug perturbation effects on gene set activity, differential Vision scores were calculated as the average difference in Vision scores between drug-treated and untreated control cells. Comparisons were performed for each plate and each cell line independently, with drug-treated cells compared against vehicle (DMSO)-treated controls within the same plate. Comparisons with fewer than 50 cells in either condition were discarded, leaving 62,927 comparisons.

For each comparison, 250 Monte Carlo samples of estimated normalized gene expression levels for all 62,710 genes were generated using the SCVI model for both drug-treated and control cells. These samples were converted into gene-set scores based on the filtered gene sets, resulting in 250 Monte Carlo samples of differential gene-set scores. The median differential gene-set scores across Monte Carlo samples were used as the final differential gene-set scores for each of the 7,390 gene sets, generating a matrix of differential gene-set scores across 62,927 comparisons.

### Dimensionality reduction of differential gene-set scores

The first 50 principal components of the differential gene-set scores of the 7,390 gene sets were used as input to the TSNE function in the cuML package from NVIDIA RAPIDS with default parameters to create a two-dimensional visualization of the comparisons.

We implemented neural network multidimensional scaling (NNMDS), a scalable solution to the metric multidimensional scaling problem (Canzar et al. 2024). Our implementation followed the original publication, with the modification of setting the width of the intermediate neural network layer to 10 times the size of the input dimensionality. The first 128 principal components of the 7,390 gene set scores were used as input to NNMDS, with a two-dimensional target for visualization. Training was performed over 250 epochs using the 62,927 comparisons.

For visualization and illustration of differential gene-set scores, comparisons from the replication plate, plate 14 (copy of plate 6), were excluded, leaving 57,471 comparisons.

### E-distance

Using the previously computed scVI 10-dimensional latent representation, E-distance metric was calculated as described in (Peidli et al. 2024). Briefly, all pairwise euclidean distances were calculated for the control (“DMSO_TF”) cell population and separately for the treated cell population. The average of the control and treated pairwise distances was then subtracted from 2x the average pairwise euclidean distances between all control and treated cells. For the final E-distance measure, this value was then adjusted for the total number of cells being considered. Within each plate, we calculated E-distance for all drug treatment conditions compared to control (DMSO_TF) for a given cell line. For cell line–drug conditions with multiple plates, a median was taken across all plates for a single aggregated E-distance metric. To then summarize the strength of a given drug, the median value for that drug was taken across all cell lines.

For comparison of E-distance metric to other datasets, namely Sciplex3 (Srivatsan et al. 2020) and the Replogle2022 genome-scale CRISPRi dataset (Replogle et al. 2022), we first downloaded datasets from the *scperturb* dataset collection (http://projects.sanderlab.org/scperturb/) provided by the Sander lab (Peidli et al. 2024). These datasets were filtered to include >50 cells per perturbation/treatment condition then run through scVI with 10-dimensional latent representation. E-distances were then calculated as described for the Tahoe-100M dataset.

### Cell cycle phase analysis

Cell cycle classification was conducted using Scanpy’s “score_genes_cell_cycle” function with gene sets sourced from the Regev lab cell cycle signature (https://github.com/scverse/scanpy_usage/blob/master/180209_cell_cycle/data/regev_lab_cell_cycle_genes.txt). Cells were assigned to the G1, S, or G2/M phase according to this signature, after which an enrichment score for each phase was calculated for every cell line, drug treatment combination. This enrichment score, defined as the log-odds ratio log2((phase_drug/nonphase_drug)/(phase_ctrl/nonphase_ctrl)), facilitated a direct comparison between cells treated with a drug and their corresponding control (DMSO_TF) counterparts. To address potential plate batch effects, the score was computed on a per cell line, per plate basis.

Subsequently, enrichment scores for each cell cycle phase were compared across different drug mechanisms of action (MOAs) to assess whether specific drug classes induced distinct alterations in cell cycle phase distributions. For this analysis drugs without clear MOAs were removed.

### Gene-set signature analysis

To assess the transcriptional impact of treatments targeting the RAS/RAF signaling pathway, we examined inhibitors that specifically modulate this pathway, including pan-RAS (RMC-6236), BRAF-V600E (Dabrafenib) inhibitors. We first filtered the dataset to retain cells treated with these inhibitors along with their corresponding plate-based controls, and further subset the expression matrix to include only protein-coding genes from MSigDB. Raw expression values were then UMI-scaled on a per-cell basis and further subset by selecting the top 50% most variable genes based on standard deviation. This generated a normalized expression matrix that was used as input for subsequent analyses with Vision (DeTomaso et al. 2019). For these analyses, KRAS-associated pathways from MSigDB were selected and filtered to include only the highly variable genes identified in our dataset. Vision was then run to derive pathway scores for these refined KRAS signatures. For each combination of cell line and drug treatment, we computed an aggregate differential Vision score using the equation (treat_mdn - dmso_mdn) / (treat_iqr + dmso_iqr), which reflects the difference between the median Vision scores for the treatment and the DMSO control, normalized by the combined interquartile range. Finally, we compared these differential Vision scores across treatments using the KRAS.600.LUNG.BREAST.V1 signature after stratifying cell lines into three groups: those harboring KRAS mutations without concurrent RAF mutations, those with BRAF V600E mutations absent RAS mutations, and those lacking both RAS and RAF mutations.

## Data Availability

Data is available on HuggingFace.

## Acknowledgments

We thank the Parse Biosciences organization for their support of this project with Gigalab services. We thank Ultima Genomics for their support of this project with sequencing services. We thank the team members at Tahoe Therapeutics for their support of the organization: Yasmin Sameni, Chancy Fessler, Shreshth Gandhi and Farnoosh Javadi. We thank Lilly Gateway Labs for their hosting of Tahoe Therapeutics’ laboratories.

## References

Adamson, Britt, Thomas M. Norman, Marco Jost, Min Y. Cho, James K. Nuñez, Yuwen Chen, Jacqueline E. Villalta, et al. 2016. “A Multiplexed Single-Cell CRISPR Screening Platform Enables Systematic Dissection of the Unfolded Protein Response.” Cell 167 (7): 1867–82.e21. 10.1016/j.cell.2016.11.048.

Aldridge, Sarah, and Sarah A. Teichmann. 2020. “Single Cell Transcriptomics Comes of Age.” Nature Communications 11 (1): 4307. 10.1038/s41467-020-18158-5.

Almogy, Gilad, Mark Pratt, Florian Oberstrass, Linda Lee, Dan Mazur, Nate Beckett, Omer Barad, et al. 2022. “Cost-Efficient Whole Genome-Sequencing Using Novel Mostly Natural Sequencing-by-Synthesis Chemistry and Open Fluidics Platform.” bioRxiv. 10.1101/2022.05.29.493900.

Beutler, Andreas S., Side Li, Rebekka Nicol, and Martin J. Walsh. 2005. “Carbamazepine Is an Inhibitor of Histone Deacetylases.” Life Sciences 76 (26): 3107–15. 10.1016/j.lfs.2005.01.003.

Bock, Christoph, Paul Datlinger, Florence Chardon, Matthew A. Coelho, Matthew B. Dong, Keith A. Lawson, Tian Lu, et al. 2022. “High-Content CRISPR Screening.” Nature Reviews Methods Primers 2 (1): 1–23. 10.1038/s43586-021-00093-4.

Buettner, Florian, Kedar N. Natarajan, F. Paolo Casale, Valentina Proserpio, Antonio Scialdone, Fabian J. Theis, Sarah A. Teichmann, John C. Marioni, and Oliver Stegle. 2015. “Computational Analysis of Cell-to-Cell Heterogeneity in Single-Cell RNA-Sequencing Data Reveals Hidden Subpopulations of Cells.” Nature Biotechnology, January. 10.1038/nbt.3102.

Bunne, Charlotte, Yusuf Roohani, Yanay Rosen, Ankit Gupta, Xikun Zhang, Marcel Roed, Theo Alexandrov, et al. 2024. “How to Build the Virtual Cell with Artificial Intelligence: Priorities and Opportunities.” Cell 187 (25): 7045–63. 10.1016/j.cell.2024.11.015.

Canzar, Stefan, Van Hoan Do, Slobodan Jelić, Sören Laue, Domagoj Matijević, and Tomislav Prusina. 2024. “Metric Multidimensional Scaling for Large Single-Cell Datasets Using Neural Networks.” Algorithms for Molecular Biology: AMB 19 (1): 21. 10.1186/s13015-024-00265-3.

Cao, Junyue, Jonathan S. Packer, Vijay Ramani, Darren A. Cusanovich, Chau Huynh, Riza Daza, Xiaojie Qiu, et al. 2017. “Comprehensive Single-Cell Transcriptional Profiling of a Multicellular Organism.” Science 357 (6352): 661–67. 10.1126/science.aam8940.

Čermák, Vladimír, Vojtěch Dostál, Michael Jelínek, Lenka Libusová, Jan Kovář, Daniel Rösel, and Jan Brábek. 2020. “Microtubule-Targeting Agents and Their Impact on Cancer Treatment.” European Journal of Cell Biology 99 (4): 151075. 10.1016/j.ejcb.2020.151075.

Chen, Junyi, Xiaoying Wang, Anjun Ma, Qi-En Wang, Bingqiang Liu, Lang Li, Dong Xu, and Qin Ma. 2022. “Deep Transfer Learning of Cancer Drug Responses by Integrating Bulk and Single-Cell RNA-Seq Data.” Nature Communications 13 (1): 6494. 10.1038/s41467-022-34277-7.

Crozier, Lisa, Reece Foy, Brandon L. Mouery, Robert H. Whitaker, Andrea Corno, Christos Spanos, Tony Ly, Jeanette Gowen Cook, and Adrian T. Saurin. 2022. “CDK4/6 Inhibitors Induce Replication Stress to Cause Long-Term Cell Cycle Withdrawal.” The EMBO Journal 41 (6): e108599. 10.15252/embj.2021108599.

CZI Cell Science Program, Shibla Abdulla, Brian Aevermann, Pedro Assis, Seve Badajoz, Sidney M. Bell, Emanuele Bezzi, et al. 2025. “CZ CELLxGENE Discover: A Single-Cell Data Platform for Scalable Exploration, Analysis and Modeling of Aggregated Data.” Nucleic Acids Research 53 (D1): D886–900. 10.1093/nar/gkae1142.

Dann, Emma, Neil C. Henderson, Sarah A. Teichmann, Michael D. Morgan, and John C. Marioni. 2022. “Differential Abundance Testing on Single-Cell Data Using K-Nearest Neighbor Graphs.” Nature Biotechnology 40 (2): 245–53. 10.1038/s41587-021-01033-z.

Datlinger, Paul, André F. Rendeiro, Thorina Boenke, Martin Senekowitsch, Thomas Krausgruber, Daniele Barreca, and Christoph Bock. 2021. “Ultra-High-Throughput Single-Cell RNA Sequencing and Perturbation Screening with Combinatorial Fluidic Indexing.” Nature Methods 18 (6): 635–42. 10.1038/s41592-021-01153-z.

DeTomaso, David, Matthew G. Jones, Meena Subramaniam, Tal Ashuach, Chun J. Ye, and Nir Yosef. 2019. “Functional Interpretation of Single Cell Similarity Maps.” Nature Communications 10 (1): 4376. 10.1038/s41467-019-12235-0.

Dixit, Atray, Oren Parnas, Biyu Li, Jenny Chen, Charles P. Fulco, Livnat Jerby-Arnon, Nemanja D. Marjanovic, et al. 2016. “Perturb-Seq: Dissecting Molecular Circuits with Scalable Single-Cell RNA Profiling of Pooled Genetic Screens.” Cell 167 (7): 1853–66.e17. 10.1016/j.cell.2016.11.038.

Dong, Mingze, Bao Wang, Jessica Wei, Antonio H. de O Fonseca, Curtis J. Perry, Alexander Frey, Feriel Ouerghi, et al. 2023. “Causal Identification of Single-Cell Experimental Perturbation Effects with CINEMA-OT.” Nature Methods 20 (11): 1769–79. 10.1038/s41592-023-02040-5.

Endy, D., and R. Brent. 2001. “Modelling Cellular Behaviour.” Nature 409 (6818): 391–95. 10.1038/35053181.

Ergen, Can, Valeh Valiollah Pour Amiri, Martin Kim, Aaron Streets, Adam Gayoso, and Nir Yosef. 2024. “Scvi-Hub: An Actionable Repository for Model-Driven Single Cell Analysis.” bioRxiv. 10.1101/2024.03.01.582887.

Gayoso, Adam, Romain Lopez, Galen Xing, Pierre Boyeau, Valeh Valiollah Pour Amiri, Justin Hong, Katherine Wu, et al. 2022. “A Python Library for Probabilistic Analysis of Single-Cell Omics Data.” Nature Biotechnology 40 (2): 163–66. 10.1038/s41587-021-01206-w.

Jinek, Martin, Krzysztof Chylinski, Ines Fonfara, Michael Hauer, Jennifer A. Doudna, and Emmanuelle Charpentier. 2012. “A Programmable Dual-RNA-Guided DNA Endonuclease in Adaptive Bacterial Immunity.” Science (New York, N.Y.) 337 (6096): 816–21. 10.1126/science.1225829.

Kang, Hyun Min, Meena Subramaniam, Sasha Targ, Michelle Nguyen, Lenka Maliskova, Elizabeth McCarthy, Eunice Wan, et al. 2018. “Multiplexed Droplet Single-Cell RNA-Sequencing Using Natural Genetic Variation.” Nature Biotechnology 36 (1): 89–94. 10.1038/nbt.4042.

Kantarjian, H. M., M. Talpaz, V. Santini, A. Murgo, B. Cheson, and S. M. O’Brien. 2001. “Homoharringtonine: History, Current Research, and Future Direction.” Cancer 92 (6): 1591–1605. 10.1002/1097-0142(20010915)92:6<1591::aid-cncr1485>3.0.co;2-u.

Korsunsky, Ilya, Nghia Millard, Jean Fan, Kamil Slowikowski, Fan Zhang, Kevin Wei, Yuriy Baglaenko, Michael Brenner, Po-Ru Loh, and Soumya Raychaudhuri. 2019. “Fast, Sensitive and Accurate Integration of Single-Cell Data with Harmony.” Nature Methods 16 (12): 1289–96. 10.1038/s41592-019-0619-0.

Li, Heng. 2011. “A Statistical Framework for SNP Calling, Mutation Discovery, Association Mapping and Population Genetical Parameter Estimation from Sequencing Data.” Bioinformatics (Oxford, England) 27 (21): 2987–93. 10.1093/bioinformatics/btr509.

Li, Zekun, Gerui Liu, Xiaoxiao Yang, Meng Shu, Wen Jin, Yang Tong, Xiaochuan Liu, Yuting Wang, Jiapei Yuan, and Yang Yang. 2024. “An Atlas of Cell-Type-Specific Interactome Networks across 44 Human Tumor Types.” Genome Medicine 16 (1): 30. 10.1186/s13073-024-01303-w.

Lopez, Romain, Jeffrey Regier, Michael B. Cole, Michael I. Jordan, and Nir Yosef. 2018. “Deep Generative Modeling for Single-Cell Transcriptomics.” Nature Methods 15 (12): 1053–58. 10.1038/s41592-018-0229-2.

Lotfollahi, Mohammad, Mohsen Naghipourfar, Malte D. Luecken, Matin Khajavi, Maren Büttner, Marco Wagenstetter, Žiga Avsec, et al. 2022. “Mapping Single-Cell Data to Reference Atlases by Transfer Learning.” Nature Biotechnology 40 (1): 121–30. 10.1038/s41587-021-01001-7.

Luecken, Malte D., M. Büttner, K. Chaichoompu, A. Danese, M. Interlandi, M. F. Mueller, D. C. Strobl, et al. 2022. “Benchmarking Atlas-Level Data Integration in Single-Cell Genomics.” Nature Methods 19 (1): 41–50. 10.1038/s41592-021-01336-8.

Macaulay, Iain C., Valentine Svensson, Charlotte Labalette, Lauren Ferreira, Fiona Hamey, Thierry Voet, Sarah A. Teichmann, and Ana Cvejic. 2016. “Single-Cell RNA-Sequencing Reveals a Continuous Spectrum of Differentiation in Hematopoietic Cells.” Cell Reports 14 (4): 966–77. 10.1016/j.celrep.2015.12.082.

Macosko, Evan Z., Anindita Basu, Rahul Satija, James Nemesh, Karthik Shekhar, Melissa Goldman, Itay Tirosh, et al. 2015. “Highly Parallel Genome-Wide Expression Profiling of Individual Cells Using Nanoliter Droplets.” Cell 161 (5): 1202–14. 10.1016/j.cell.2015.05.002.

McFarland, James M., Brenton R. Paolella, Allison Warren, Kathryn Geiger-Schuller, Tsukasa Shibue, Michael Rothberg, Olena Kuksenko, et al. 2020. “Multiplexed Single-Cell Transcriptional Response Profiling to Define Cancer Vulnerabilities and Therapeutic Mechanism of Action.” Nature Communications 11 (1): 4296. 10.1038/s41467-020-17440-w.

Mitchell, Jana M., James Nemesh, Sulagna Ghosh, Robert E. Handsaker, Curtis J. Mello, Daniel Meyer, Kavya Raghunathan, et al. 2020. “Mapping Genetic Effects on Cellular Phenotypes with ‘cell Villages.’” bioRxiv. bioRxiv. 10.1101/2020.06.29.174383.

Papalexi, Efthymia, Eleni P. Mimitou, Andrew W. Butler, Samantha Foster, Bernadette Bracken, William M. Mauck 3rd, Hans-Hermann Wessels, et al. 2021. “Characterizing the Molecular Regulation of Inhibitory Immune Checkpoints with Multimodal Single-Cell Screens.” Nature Genetics 53 (3): 322–31. 10.1038/s41588-021-00778-2.

Parry, David, Timothy Guzi, Frances Shanahan, Nicole Davis, Deepa Prabhavalkar, Derek Wiswell, Wolfgang Seghezzi, et al. 2010. “Dinaciclib (SCH 727965), a Novel and Potent Cyclin-Dependent Kinase Inhibitor.” Molecular Cancer Therapeutics 9 (8): 2344–53. 10.1158/1535-7163.MCT-10-0324.

Pe’er, Dana. 2005. “Bayesian Network Analysis of Signaling Networks: A Primer.” Science’s STKE: Signal Transduction Knowledge Environment 2005 (281): l4. 10.1126/stke.2812005pl4.

Peidli, Stefan, Tessa D. Green, Ciyue Shen, Torsten Gross, Joseph Min, Samuele Garda, Bo Yuan, et al. 2024. “scPerturb: Harmonized Single-Cell Perturbation Data.” Nature Methods 21 (3): 531–40. 10.1038/s41592-023-02144-y.

Rajput, Sandeep, Nimmish Khera, Zhanfang Guo, Jeremy Hoog, Shunqiang Li, and Cynthia X. Ma. 2016. “Inhibition of Cyclin Dependent Kinase 9 by Dinaciclib Suppresses Cyclin B1 Expression and Tumor Growth in Triple Negative Breast Cancer.” Oncotarget 7 (35): 56864–75. 10.18632/oncotarget.10870.

Replogle, Joseph M., Reuben A. Saunders, Angela N. Pogson, Jeffrey A. Hussmann, Alexander Lenail, Alina Guna, Lauren Mascibroda, et al. 2022. “Mapping Information-Rich Genotype-Phenotype Landscapes with Genome-Scale Perturb-Seq.” Cell 185 (14): 2559–75.e28. 10.1016/j.cell.2022.05.013.

Rosenberg, Alexander B., Charles M. Roco, Richard A. Muscat, Anna Kuchina, Paul Sample, Zizhen Yao, Lucas T. Graybuck, et al. 2018. “Single-Cell Profiling of the Developing Mouse Brain and Spinal Cord with Split-Pool Barcoding.” Science 360 (6385): 176–82. 10.1126/science.aam8999.

Schiebinger, Geoffrey, Jian Shu, Marcin Tabaka, Brian Cleary, Vidya Subramanian, Aryeh Solomon, Joshua Gould, et al. 2019. “Optimal-Transport Analysis of Single-Cell Gene Expression Identifies Developmental Trajectories in Reprogramming.” Cell 176 (6): 1517. 10.1016/j.cell.2019.02.026.

Shalem, Ophir, Neville E. Sanjana, and Feng Zhang. 2015. “High-Throughput Functional Genomics Using CRISPR-Cas9.” Nature Reviews. Genetics 16 (5): 299–311. 10.1038/nrg3899.

Simmons, Sean K., Gila Lithwick-Yanai, Xian Adiconis, Florian Oberstrass, Nika Iremadze, Kathryn Geiger-Schuller, Pratiksha I. Thakore, et al. 2023. “Mostly Natural Sequencing-by-Synthesis for scRNA-Seq Using Ultima Sequencing.” Nature Biotechnology 41 (2): 204–11. 10.1038/s41587-022-01452-6.

Skinnider, Michael A., Jordan W. Squair, Claudia Kathe, Mark A. Anderson, Matthieu Gautier, Kaya J. E. Matson, Marco Milano, et al. 2021. “Cell Type Prioritization in Single-Cell Data.” Nature Biotechnology 39 (1): 30–34. 10.1038/s41587-020-0605-1.

Squair, Jordan W., Matthieu Gautier, Claudia Kathe, Mark A. Anderson, Nicholas D. James, Thomas H. Hutson, Rémi Hudelle, et al. 2021. “Confronting False Discoveries in Single-Cell Differential Expression.” Nature Communications 12 (1): 5692. 10.1038/s41467-021-25960-2.

Srivatsan, Sanjay R., José L. McFaline-Figueroa, Vijay Ramani, Lauren Saunders, Junyue Cao, Jonathan Packer, Hannah A. Pliner, et al. 2020. “Massively Multiplex Chemical Transcriptomics at Single-Cell Resolution.” Science (New York, N.Y.) 367 (6473): 45–51. 10.1126/science.aax6234.

Subramanian, Aravind, Pablo Tamayo, Vamsi K. Mootha, Sayan Mukherjee, Benjamin L. Ebert, Michael A. Gillette, Amanda Paulovich, et al. 2005. “Gene Set Enrichment Analysis: A Knowledge-Based Approach for Interpreting Genome-Wide Expression Profiles.” Proceedings of the National Academy of Sciences of the United States of America 102 (43): 15545–50. 10.1073/pnas.0506580102.

Trapnell, Cole, Davide Cacchiarelli, Jonna Grimsby, Prapti Pokharel, Shuqiang Li, Michael Morse, Niall J. Lennon, Kenneth J. Livak, Tarjei S. Mikkelsen, and John L. Rinn. 2014. “The Dynamics and Regulators of Cell Fate Decisions Are Revealed by Pseudotemporal Ordering of Single Cells.” Nature Biotechnology 32 (4): 381–86. 10.1038/nbt.2859.

Tsherniak, Aviad, Francisca Vazquez, Phil G. Montgomery, Barbara A. Weir, Gregory Kryukov, Glenn S. Cowley, Stanley Gill, et al. 2017. “Defining a Cancer Dependency Map.” Cell 170 (3): 564–76.e16. 10.1016/j.cell.2017.06.010.

Tyler, Michael, Avishai Gavish, Chaya Barbolin, Roi Tschernichovsky, Rouven Hoefflin, Michael Mints, Sidharth V. Puram, and Itay Tirosh. 2024. “The Curated Cancer Cell Atlas: Comprehensive Characterisation of Tumours at Single-Cell Resolution.” bioRxiv. 10.1101/2024.10.11.617836.

Virshup, Isaac, Sergei Rybakov, Fabian J. Theis, Philipp Angerer, and F. Alexander Wolf. 2021. “Anndata: Annotated Data.” bioRxiv. 10.1101/2021.12.16.473007.

Weinberger, Ethan, Chris Lin, and Su-In Lee. 2023. “Isolating Salient Variations of Interest in Single-Cell Data with contrastiveVI.” Nature Methods 20 (9): 1336–45. 10.1038/s41592-023-01955-3.

Xie, Yi, Huimei Chen, Vasuki Ranjani Chellamuthu, Ahmad Bin Mohamed Lajam, Salvatore Albani, Andrea Hsiu Ling Low, Enrico Petretto, and Jacques Behmoaras. 2024. “Comparative Analysis of Single-Cell RNA Sequencing Methods with and without Sample Multiplexing.” International Journal of Molecular Sciences 25 (7). 10.3390/ijms25073828.

Yu, Channing, Aristotle M. Mannan, Griselda Metta Yvone, Kenneth N. Ross, Yan-Ling Zhang, Melissa A. Marton, Bradley R. Taylor, et al. 2016. “High-Throughput Identification of Genotype-Specific Cancer Vulnerabilities in Mixtures of Barcoded Tumor Cell Lines.” Nature Biotechnology 34 (4): 419–23. 10.1038/nbt.3460.

Yu, Johnny X., Jung Min Suh, Katerina D. Popova, Kristle Garcia, Tanvi Joshi, Bruce Culbertson, Jessica B. Spinelli, et al. 2024. “Multiplexed Mosaic Tumor Models Reveal Natural Phenotypic Variations in Drug Response within and between Populations.” bioRxiv. 10.1101/2024.12.13.628239.

Zhong, Mengya, Jinshui Tan, Guangchao Pan, Yuelong Jiang, Hui Zhou, Qian Lai, Qinwei Chen, et al. 2021. “Preclinical Evaluation of the HDAC Inhibitor Chidamide in Transformed Follicular Lymphoma.” Frontiers in Oncology 11 (December):780118. 10.3389/fonc.2021.780118.

